# Unlocking Precision Diagnostics: A Multimodal Framework Integrating Metabolomics with Advanced Machine Learning Techniques

**DOI:** 10.1101/2025.01.19.633800

**Authors:** Parisa Shahnazari, Kaveh Kavousi, Hamid Reza Khorram Khorshid, Bahram Goliaei, Reza M Salek

## Abstract

Integrating multiple omics modalities is a crucial strategy in cancer research, particularly in metabolomics, enabling early detection and detailed exploration of cancer biomarker signatures. This study evaluates five strategies for integrating metabolomics data from liquid chromatography-mass spectrometry, gas chromatography-mass spectrometry, and nuclear magnetic resonance. Deep Transfer Learning and Multiple Kernel Learning demonstrated superior performance, significantly improving classification accuracy, sensitivity, and robustness compared to single-modality analyses. Deep Transfer Learning employed a custom autoencoder for feature extraction followed by artificial neural network classification, while Multiple Kernel Learning optimized kernel matrices across different modalities. Feature extraction in the Deep Transfer Learning approach, combined with the selection of important features and subsequent analysis, revealed elevated levels of monounsaturated phospholipids such as phosphatidylcholine 30:1, phosphatidylethanolamine 32:1, and sphingomyelin 32:1 in HER2-positive cases. Additionally, β-alanine, gluconic acid, and N-acetylaspartic acid were increased, whereas 5’-deoxy-5’-methylthioadenosine and nicotinamide were decreased. These methods advance cancer detection, biomarker discovery, and the development of precise diagnostic and therapeutic tools while offering robust and adaptable strategies for multi-omics data integration across diverse biological datasets.

## 1 Introduction

Metabolomics, the comprehensive study of small molecules within biological systems, holds immense potential for elucidating the intricate molecular metabolic profiles associated with human diseases (1). However, comprehensively detecting all metabolites using a single analytical technology presents significant challenges due to their diverse chemical nature and the limitations of current analytical instruments. As a result, a typical metabolomics experiment using a single analytical technique can only provide a partial identification or discovery of the full spectrum of metabolites within a system. Common analytical platforms employed in metabolomic investigations include liquid chromatography-mass spectrometry (LC-MS), gas chromatography-mass spectrometry (GC-MS), and nuclear magnetic resonance spectroscopy (NMR) (2). Holistic coverage of the metabolome can be achieved by integrating NMR, GC-MS, and LC-MS for comprehensive analysis. Each platform captures a unique set of metabolites, collectively offering a more detailed perspective (2, 3). NMR detects high-concentration compounds with limited sensitivity, GC-MS is ideally suited for volatile compounds, and LC-MS is versatile in covering a broad spectrum of compounds and concentrations with high sensitivity (4). *Fatema Bhinderwala* and colleagues (2018) highlighted the benefits of combining mass spectrometry and NMR for improved detection and annotation(5).

Multi-modal integration, or multi-view, multi-platform, or cross-modal integration, is a powerful strategy that significantly reduces false positives and negatives (6). By implementing integration techniques, analytical confidence is notably bolstered as results are validated across platforms, thus enhancing the reliability and robustness of the data(7). Moreover, integrating different platforms amplifies the likelihood of discovering previously unidentified metabolites through cross-referencing various spectra or analytes, effectively resolving ambiguities in compound identification(1). In addition, comprehensive coverage in metabolomic studies contributes to a more profound understanding of metabolic pathways, enabling a more detailed pathway analysis (6). Integrating multimodal metabolomics data is a valuable strategy in cancer research. It helps identify early detection biomarkers and monitor perturbations in metabolic profiles in biological fluids and tissues before clinical symptoms manifest. This proactive approach facilitates early disease diagnosis and is crucial in predicting clinical outcomes, advancing personalized medicine, and drug discovery (1, 8, 9).

On the other hand, integrating multiple platforms can be complex and challenging. This process requires careful consideration due to resource-intensive aspects that involve intricate sample preparation procedures, substantial time commitments, and the need for specialized expertise. Addressing these challenges is essential to ensure reproducibility, facilitate a smooth integration process, and derive meaningful interpretations from the diverse analytical techniques. Integration methodologies can be broadly categorized as either graph-based or non-graph-based, depending on the characteristics of the datasets and the functionalities of techniques employed. *Eugene Lin* et al. (2017) investigated the integration of multimodals in omics by applying machine learning and systems genomics approaches (10). *Milan Picard* et al. (2021) categorized five integration strategies—early, mixed, intermediate, late, and hierarchical. This classification also involved categorizing methods based on mathematical aspects, such as network-based, deep learning-based, kernel-based, and matrix factorization-based approaches for integrating multi-omics data(8). *Yanlin Wang* et al. (2022) have further ascribed the concept of intermediate integration by categorizing it based on matrix factorization, component analysis, multiple kernel learning, similarity network, Bayesian Network, Graph Transformation, and Artificial Neural Networks (11). Nasim Vahabi *et al.* (2022) categorized unsupervised multi-omics data integration into three comprehensive categories: Regression/Association-based, Clustering, and Network-based Methods(12). Additionally, *Francis E. Agamah* (2022) investigated network-based integrative multiomics(13). Furthermore, *Daniyal Rajput* et al. (2023) observed a stabilization in both effect sizes and ML accuracies beyond a certain sample threshold(14). This finding suggests that even a modest sample size can suffice for analysis when dealing with high-quality datasets. The study proposed criteria for determining the appropriate sample size, emphasizing the importance of substantial effect sizes (≥ 0.5) and a high machine-learning accuracy threshold (≥ 80%). This recommendation is particularly valuable for metabolomics data, where the number of samples is typically limited.

This study evaluates five integration methodologies for multimodal data to identify the most effective approaches in metabolomics. The methods include straightforward concatenation, concatenated-ensemble, deep forest, multiple kernel learning, and deep transfer learning. In the deep transfer learning approach, features extracted from the encoder are weighted, and key features are selected for further analysis to identify significant biomarker signatures.

The primary objective of this research is to develop a robust pipeline applicable to medicine, enhancing the precision and depth of insights provided by analytical models. The integration techniques employed in this study can be summarized as follows:

### Straightforward Concatenation

Straightforward concatenation is a common method for horizontally combining datasets with identical samples and class labels, frequently used in biological research to achieve more comprehensive insights. By expanding the dataset’s feature space, this approach enhances pattern recognition through the integration of diverse platform data. However, it also presents challenges such as increased dimensionality, potential redundancy, platform compatibility assumptions, and data quality and interpretability concerns. While feature selection can help reduce dimensionality and redundancy, it cannot fully address platform compatibility and data quality issues. Additional preprocessing steps, such as normalization or alignment of datasets, are often required to ensure compatibility. Furthermore, this fusion technique typically serves as an initial step for integrating advanced machine learning methods, further enhancing the dataset’s analytical power and interpretability (15–17).

### Concatenation-Ensemble

The Concatenation-Ensemble method employs ensemble learning, which integrates multiple learning algorithms to enhance predictive performance beyond that of individual models. This approach involves the horizontal integration of datasets from diverse sources, followed by applying ensemble techniques such as Stacking, Bagging, AdaBoost, or Voting on the consolidated dataset. To ensure the robustness and reliability of the model, cross-validation is employed, validating its performance across various data subsets. By leveraging the strengths of multiple models and compensating for their weaknesses, this strategy significantly improves overall predictive accuracy and generalization capabilities(15).

### Deep Forest

Deep Forest is an advanced ensemble learning technique introduced by Zhou and Feng in 2017, designed for both classification and regression tasks(18). It leverages multiple layers of tree-based models, typically random forests, in a cascading architecture where each layer processes the input data and the output from the previous layer, including class probabilities or regression estimates. This hierarchical structure allows the model to represent features at different levels of detail, effectively capturing complex data patterns. The process starts with an initial layer that applies tree-based models to the original input features. Each subsequent layer then receives a combination of the original features and the outputs from the previous layer, refining the data representation and enhancing predictive accuracy(19, 20).

### Multiple Kernel Learning

Multiple Kernel Learning (MKL) is a robust paradigm that addresses diverse learning challenges, particularly in classification and regression. Jérôme Mariette et al. (2017) and Xingheng Yu *et al*. (2020) underscored MKL’s efficacy when applied to breast cancer heterogeneous data, demonstrating commendable performance outcomes(21, 22). The essence of MKL lies in its ability to seamlessly integrate information from multiple sources, as reflected in the combined kernel function k (x, x’):

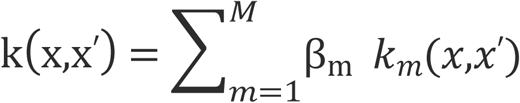

where k (x, x’) characterizes the combined kernel function between two input data points, x, and x,’ considering information from M distinct platforms.

The summation 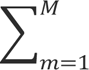 denotes the weighted sum of individual kernel functions. km (x, x’), denotes the individual kernel function associated with the m-th platform. *β*_*m*_ represents the weight or coefficient assigned to the m-th individual kernel (23). The weights *β*_*m*_ play a crucial rol in adjusting the influence of each platform on the combined kernel.

### Deep Transfer Learning

Transfer learning is an advanced machine learning technique that employs knowledge gained from one task or dataset to enhance performance on a related task or different dataset. Unlike traditional models trained independently for each task, transfer learning draws on insights from earlier tasks to improve generalization and adaptability in new situations. This method is particularly beneficial in scenarios with limited data, allowing for more effective learning by utilizing established features and patterns(24).

Conventional transfer learning modifies models for related tasks by integrating knowledge from one domain to improve performance in another. In contrast, concatenation transfer learning combines features from multiple sources before feeding them into the model, providing a more comprehensive understanding when each source contributes unique information (25).

This study applies transfer learning using a custom autoencoder architecture consisting of an encoder and a bottleneck (hidden layer). The encoder compresses input data into a lower-dimensional representation, capturing the most essential features through the bottleneck. However, unlike traditional autoencoders, the decoder is not used for data reconstruction. Instead, the bottleneck layer serves as the key feature extraction point, condensing data into a compact, informative form. In addition, a pre-trained encoder, previously trained on a related task, is employed to capture general patterns and feature representations. This pre-training provides a strong initialization, enabling the transfer of learned features to the new task, and improving performance, especially on limited datasets, by reducing the need for extensive re-training and speeding up convergence (26, 27).

### SHapley Additive exPlanations (SHAP)

SHAP (SHapley Additive exPlanations) is a unified framework for interpreting machine learning model predictions based on game theory’s Shapley values (28). It calculates each feature’s contribution to a prediction by comparing predictions with and without that feature across different combinations of other features. SHAP helps identify important features by assigning importance values representing their average marginal contribution to model predictions (29, 30). Key advantages include its strong theoretical foundation with properties like local accuracy and consistency, its ability to work with any machine learning model (model-agnostic), and its provision of global and local feature importance interpretations. In cancer research, SHAP helps identify key biomarkers influencing tumor development and treatment response, enabling more precise patient stratification(31). By analyzing their unique molecular and clinical profiles in precision medicine, SHAP helps explain why specific treatments might work better for certain patients (32, 33).

## 2 Method details

### 2-1 Metabolomics multimodal datasets

The datasets utilized in this study originated from breast cancer tissue data acquired earlier as part of the METAcancer FP7 project. The metabolomic datasets were obtained from three distinct platforms: High-Resolution Magic Angle Spinning Nuclear Magnetic Resonance (HRMAS 1HNMR), Gas Chromatography-Time-of-Flight Mass Spectrometry (GC-TOFMS), and Acquity Ultra Performance LC (UPLC). For detailed information regarding the experimental setup, samples, and other related metadata, please refer to the cited references(34–37)

### 2-2 Data curation

Three distinct metabolomics platforms—NMR, GC-MS, and LC-MS—were used to profile metabolites within the HER2 group. Although earlier efforts under the METAcancer FP7 project did not succeed in achieving or reporting classification within the HER2 group, this study aimed to address these challenges. To optimize integration and address dimensionality challenges, the original datasets underwent refinement. This process aimed to ensure uniformity in the number of samples across all three platforms while preserving the unique features characteristic of each platform. A carefully curated cohort of 253 samples was established, ensuring consistent representation across all platforms. The number of features varied by platform, with 180 features for NMR, 161 for GC-MS, and 183 for LC-MS positive data. LC-MS negative datasets were excluded from this study due to a limited number of available features (34 known metabolites). The resulting harmonized dataset served as the foundation for subsequent analyses.

### 2-3 Machine Learning Framework and Implementation

All machine learning analyses were conducted using Python 3.6. The methodologies outlined below are elaborated upon and contextualized with pertinent references to enhance understanding and credibility.

### 2-4 Oversampling

Considering the significant disparity in the distribution of HER2 positive (0.1462) and HER2 negative (0.8538) instances across all three platforms, oversampling was implemented using the SMOTE (Synthetic Minority Over-sampling Technique) library in Python to rectify this discrepancy. This strategy indirectly augmented the sample sizes, yielding 423 instances for each platform, thus promoting a more balanced dataset(38).

### 2-5 Feature Selection

The differential abundance of HER2 positive versus HER2 negative instances across all three platforms did not present significant disparities compared to Cancerous versus Non-cancerous, making differential expression feature selection methods such as fold change and effect size unsuitable for this study (see supplement). Consequently, four algorithmic feature selection techniques—ReliefF, SVM-RFE, RF-RFE, and Mutual Information—were evaluated and compared to determine the optimal method for feature elimination. A brief overview of each method is provided below:

**ReliefF (Relief Feature Selection):** ReliefF is a filter-based feature selection method that gauges feature relevance by assessing their ability to distinguish instances of the same and different classes. It assigns weights to features based on their impact on classification accuracy (39).

**SVM-RFE (Support Vector Machine Recursive Feature Elimination):** SVM-RFE is a wrapper-based feature selection technique employing Support Vector Machines (SVMs) in a recursive elimination process. It iteratively removes the least important features, ranking them based on their contribution to SVM performance(40).

**RF-RFE (Random Forest Recursive Feature Elimination):** RF-RFE is a wrapper-based feature selection method utilizing Random Forests. It recursively eliminates less important features by assessing their impact on the accuracy of a Random Forest classifier. Features are ranked based on their contribution to model performance(41).

**Mutual Information:** Mutual Information is a filter-based feature selection measure quantifying the dependency between two variables. It assesses the information shared between each feature and the target variable, ranking features based on their information content and relevance(42).

The effectiveness of feature selection methods was evaluated using the SVM-Linear model with C=1. The optimal number of features for each method on individual platforms was determined. In this study, stability measures, such as the Jaccard score, were employed to assess the consistency of feature selection methods across various data subsets. The Jaccard score quantifies the similarity among selected features across different subsets. A higher Jaccard score indicates greater stability, signifying that the feature selection method is more reliable.

### 2-6 Individual platform classification

The feature selection method employed consistently across all platforms was SVM-RFE. For binary classification within the HER2 group, three SVM models—SVM-linear, SVM-RBF, and SVM-poly—were utilized. Each dataset underwent partitioning into training (80%) and testing (20%) sets, treated as independent data subsets. Repeated stratified cross-validation was employed to enhance model robustness, accompanied by a grid search for hyperparameter optimization. To mitigate overfitting, the range of C values was restricted between 0.01 and 20. Evaluation metrics for individual and integrated multimodal models encompassed accuracy, F1 score, balanced accuracy, sensitivity, specificity, and MCC. These metrics were computed using the test set as independent data. MCC proved particularly valuable in imbalanced datasets, offering a comprehensive assessment of model performance by considering false positives and negatives.

### 2-7 Integrated Multimodal Techniques

Several methodologies, including basic concatenation ensemble, multiple kernel learning (MKL), and deep transfer learning, were used to integrate multimodal data.

#### 2-7-1 Straightforward Concatenation and Concatenation Ensemble Methods

Three distinct metabolomics platforms were horizontally concatenated, aligning samples in rows and featuring distinct datasets as columns. Individual platform features underwent selection using SVM-RFE. Each dataset was scaled, normalized, and oversampled before concatenation. For the concatenation-ensemble method, base model training involved a grid search for hyperparameter tuning of the SVM polynomial kernel after splitting the data. Optimal hyperparameters were identified, including the regularization parameter (C) and polynomial degree. Subsequently, a Stacking Classifier was created, combining SVM polynomial kernel, Random Forest (RF), and XGBoost (XGB) models. In the Bagging Classifier, an RF bagging classifier was the base estimator. Logistic Regression was employed as the final estimator/meta-learning in the stacking model, introducing L2 regularization (penalty=’l2’, C=1.0) and implementing 5-fold cross-validation during the stacking process.

The study employed nested repeated stratified cross-validation to evaluate both concatenated and concatenation-ensemble models. This specialized approach ensured robust hyperparameter tuning and effective management of model complexity tailored to the challenges posed by these architectures.

#### 2-7-2 Cascade Deep Forest

In this study, the Deep Forest model was employed to classify and evaluate data from three distinct metabolomics platforms, each of which had undergone feature selection. These platforms were horizontally concatenated, aligning samples in rows and representing different datasets as columns. The Deep Forest model utilized a Cascade Forest framework, composed of multiple layers of forest estimators, including both Random Forest and Extra Trees classifiers. This hierarchical structure enabled the progressive refinement of feature representations across layers. Hyperparameter tuning focused on key parameters such as the number of layers, forests per layer, and estimators per forest.

Initially, we experimented with a multi-grained scanning approach using a sliding window. However, this led to higher risks of overfitting and lower evaluation metrics. Consequently, the multi-grained scanning was excluded from the final workflow.

Model evaluation and hyperparameter optimization were conducted using repeated stratified cross-validation. This approach ensured a robust and reliable framework for the classification and analysis of complex metabolomics datasets.

#### 2-7-3 Multiple Kernel Learning

EasyMKL, a scalable Multiple Kernel Learning (MKL) algorithm, was adeptly employed for integrating information from multiple platforms. The algorithm’s scalability and computational efficiency enable effective management of many kernels (43, 44). To optimize the MKL framework, the ‘cvxopt’ solver, renowned as a quadratic programming solver, was utilized to identify the optimal combination of kernels. The combined kernel matrix was split into training and testing sets, employing 5-fold repeated stratified cross-validation. Hyperparameter tuning was conducted for ‘lam’ and ‘C’ using grid search during the MKL process. ‘lam’ typically represents the regularization parameter controlling the trade-off between model complexity and accuracy. At the same time, ‘C’ is associated with the cost parameter, influencing the penalty for misclassification.

#### 2-7-4 Deep Transfer Learning / Multimodal Deep Learning

The autoencoders employ a carefully designed architecture featuring customized encoding dimensions, ‘Tanh’ activation functions, and optimization using the Adam optimizer with mean squared error, which has proven pivotal (36).

This approach sets the stage for feature extraction from three distinct platforms. Features from these platforms are concatenated and then partitioned into a 70% training and 30% testing split. Similarly, the ANN adopts a comparable architecture, utilizing the ‘tanh’ activation function and the Adam optimizer. Autoencoders and ANN undergo fine hyperparameter tuning to mitigate overfitting and bolster performance. This tuning is facilitated by a repeated stratified cross-validation K-Fold approach with five splits, ensuring thorough model evaluation and decreasing the risks of overfitting.

Hyperparameters are optimized using GridSearchCV from scikit-learn. This method samples hyperparameters from predefined distributions, and model performance is evaluated through cross-validation. Additionally, Gaussian noise is added to the training data to reduce overfitting, and models are reinitialized during cross-validation to ensure robustness.

Within the autoencoder framework, hyperparameter optimization encompasses various facets:

- Encoding dimension: Governing the size of the bottleneck layer and compression levels.
- Batch size: Influencing the speed of convergence and memory utilization.
- Dropout rate: A regularization technique aimed at mitigating overfitting.
- L1 and L2 penalties: Regularization mechanisms designed to discourage large parameter values.
- Learning rate: Pivotal for adjusting weights during optimizer training.

In the ANN domain, optimization extends beyond basic parameters like batch size, dropout rate, and learning rate. It includes ‘alpha’, which determines the strength of L2 regularization, thereby shaping the model’s capability to curb overfitting by penalizing large weights. the number of artificial neurons directly influencing the model’s aptitude to grasp intricate patterns.

#### 2-7-5 Feature weights, Feature Mapping, and Selecting Important Features

The process begins by aligning feature names to match the input dimensions of the encoder’s bottleneck layer weights, ensuring consistency in naming. Next, feature mapping is performed by extracting the weights of the bottleneck layer and identifying the top contributing input features for each encoded feature based on the absolute values of their weights. This creates a mapping of encoded features to their most influential input features and weights. Using this mapping, the importance of the global feature is calculated by aggregating the absolute weights of each input feature across all encoded features. The aggregated values are then ranked to identify the most essential features globally, providing insights into feature significance for individual platforms (45, 46).

#### 2-7-6 SHapley Additive exPlanations (SHAP) Analysis

The process begins with splitting the dataset. Features are then scaled to ensure uniformity in input data. Next, a **Random Forest** model is optimized through hyperparameter tuning using GridSearchCV, and the best model is selected. Feature selection is performed using Recursive Feature Elimination (RFE) to identify the most relevant features for the model. The refined model is retrained on these selected features. Subsequently, **SHAP** values are computed to quantify feature contributions for predictions, separated by classes (e.g., HER2+ and HER2-). Mean absolute SHAP values are calculated for each feature per class, enabling comparison of feature contributions. A data frame is created to rank features by importance and visualize the top contributors.

## 3 Results

### 3-1 Feature Selection

Four different methodologies were employed and evaluated to determine the most efficient feature selection method for the HER2 subtype group across various platforms: ReliefF, SVM-RFE, RF-RFE, and Mutual Information. For the assessment, SVM-Linear was used without any hyperparameter tuning (**Table 1**). The supporting information (**S1 Fig**) provides the number of optimized features for each method on individual platforms.

**Table 1:**
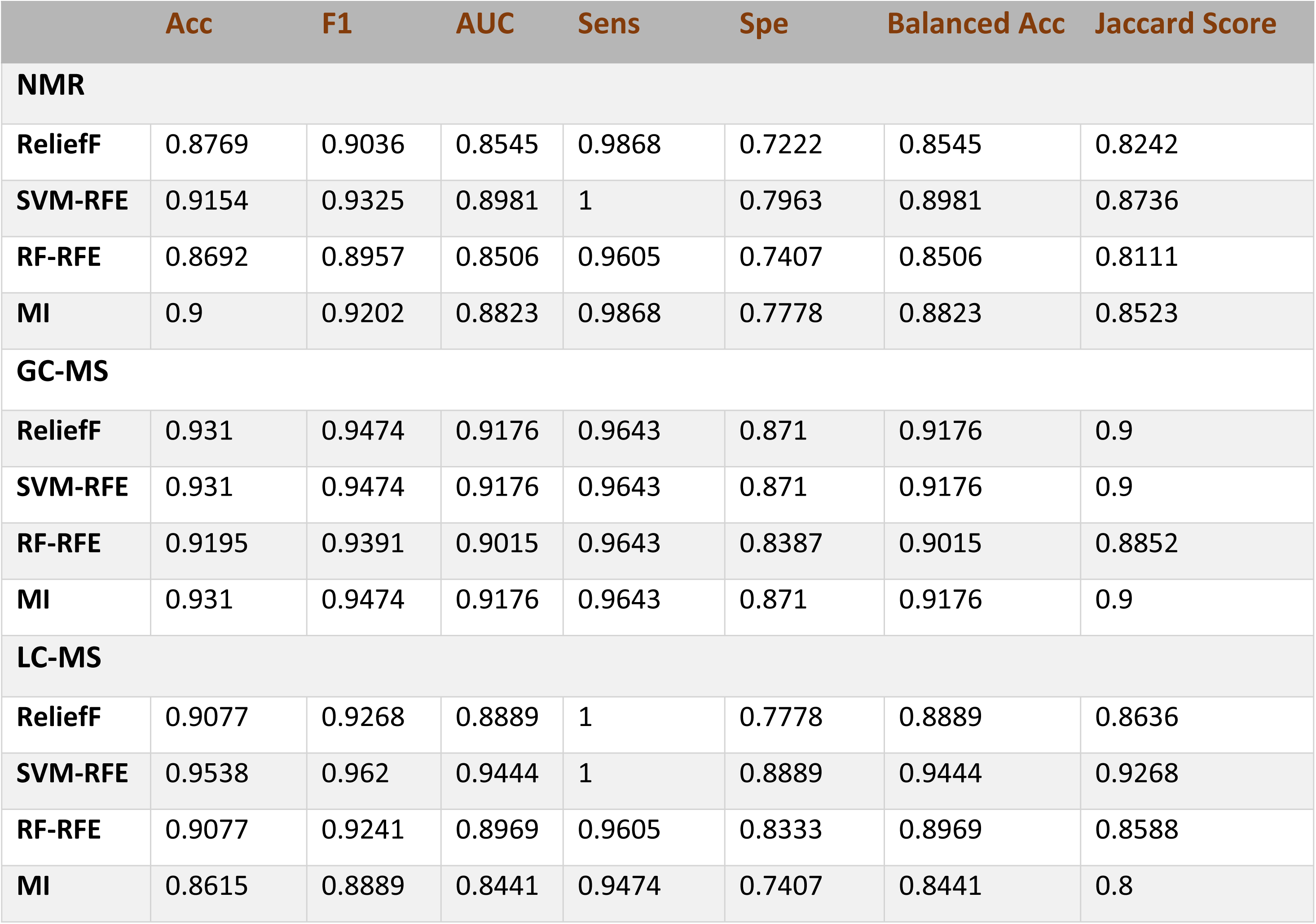
The evaluation metrics for four different feature selection methods using SVM-linear. The Jaccard score was used to assess feature stability. This score quantifies the similarity among selected features across data subsets. A higher Jaccard score indicates greater stability, meaning the feature selection is more reliable.

### 3-2 Individual Modal Classification

The evaluation metrics for three models—SVM-Linear, SVM-RBF, and SVM-Polynomial— across individual platforms are presented in **Table 2**. In the NMR dataset, SVM-Linear demonstrates superior performance. Conversely, SVM-RBF outperforms Accuracy, F1 Score, and Specificity in the GC-MS dataset. However, SVM-Linear exhibits higher metrics in Balanced Accuracy, AUC, and MCC, showcasing its overall effectiveness compared to other methods. In the LC-MS dataset, SVM-Polynomial demonstrates superior performance, characterized by higher metrics and lower standard deviations, indicating heightened robustness.

**Table 2:** A Comparison of Evaluation Metrics for Individual Metabolomic Platforms. Three SVM models—SVM-Linear, SVM-RBF, and SVM-Polynomial—were applied to each platform after feature elimination using SVM-RFE. The evaluation metrics include Accuracy (Acc), F1 Score (F1), AUC Score (AUC), Sensitivity (Sen), Specificity (Spe), Balanced Accuracy (BA), and Matthews Correlation Coefficient (MCC).

|  | Acc | F1 Score | AUC | Sen | Spe | BA | MCC |
| --- | --- | --- | --- | --- | --- | --- | --- |
| **NMR** |  |  |  |  |  |  |  |
| **SVM-Linear** | **0.9537** +/-<br>**0.0342** | **0.9533** +/-<br>**0.0334** | **0.9566** +/-<br>**0.0322** | **0.9969** +/-<br>**0.0185** | **0.9164**<br>+/- 0.0622 | **0.9566**<br>+/- 0.0322 | **0.9130** +/-<br>**0.0622** |
| **SVM-RBF** | 0.9343 +/-<br>0.0107 | 0.9512 +/-<br>0.0078 | 0.9100+/-<br>0.0136 | 0.9943<br>+/- 0.0107 | 0.8258<br>+/- 0.0274 | 0.100<br>+/- 0.0136 | 0.8591<br>+/- 0.0234 |
| **SVM-Poly** | 0.9167 +/-<br>0.0218 | 0.9340 +/-<br>0.0180 | 0.9156 +/-<br>0.0195 | 0.9193 +/-<br>0.0304 | 0.9119<br>+/- 0.0208 | 0.9156<br>+/- 0.0195 | 0.8225<br>+/- 0.0447 |
| **GC-MS** |  |  |  |  |  |  |  |
| **SVM-Linear** | 0.9429 +/-<br>0.0366 | 0.9431 +/-<br>0.0348 | **0.9466** +/-<br>**0.0345** | **0.9988**<br>+/- 0.0124 | 0.8944<br>+/- 0.0659 | **0.9466**<br>+/- **0.0345** | **0.8935**<br>+/- **0.0658** |
| **SVM-RBF** | **0.9457** +/-<br>**0.0076** | **0.9580** +/-<br>**0.0059** | 0.9394 +/-<br>0.0093 | 0.9614<br>+/- 0.0086 | **0.9174**<br>+/- **0.0184** | 0.9394<br>+/- 0.0093 | 0.8816<br>+/- 0.0167 |
| **SVM-Poly** | 0.9444 +/-<br>0.0183 | 0.9518 +/-<br>0.0137 | 0.9287 +/-<br>0.0218 | 0.9832<br>+/- 0.0159 | 0.8742<br>+/- 0.0322 | 0.9287<br>+/- 0.0218 | 0.8790<br>+/- 0.0404 |

| LC-MS |  |  |  |  |  |  |  |
| --- | --- | --- | --- | --- | --- | --- | --- |
| **SVM-Linear** | 0.9588 +/-<br>0.0319 | 0.9585 +/-<br>0.0307 | 0.9615 +/-<br>0.0298 | 1.0000<br>+/- 0.0000 | 0.9231<br>+/- 0.0569 | 0.9615<br>+/- 0.0298 | 0.9225<br>+/- 0.0576 |
| **SVM-RBF** | 0.9601 +/-<br>0.0121 | 0.9691 +/-<br>0.0094 | 0.9557 +/-<br>0.0141 | 0.9711<br>+/- 0.0128 | 0.9403<br>+/- 0.0256 | 0.9557<br>+/- 0.0147 | 0.9132<br>+/- 0.0263 |
| **SVM-Poly** | **0.9708 +/-**<br>**0.0166** | **0.9652 +/-**<br>**0.0129** | **0.9652 +/-**<br>**0.0176** | **0.9848**<br>**+/- 0.0181** | **0.9455**<br>**+/- 0.0261** | **0.9652**<br>**+/- 0.0176** | **0.9366**<br>**+/- 0.0363** |

### 3-3 Multimodal Data Integration

The study examined various multimodal techniques, including Straightforward Concatenation, Concatenation-Assembled, Deep Forest, Multiple Kernel Learning, and Deep Transfer Learning. The evaluation employed datasets, namely NMR, GC-MS, and LC-MS, which featured uniform sample quantities (matching rows) but differed in the number of features (distinct columns).

#### 3-3-1 Straightforward Concatenation

In this approach, datasets were horizontally concatenated while ensuring the preservation of identical label groups, such as HER2 positive and negative, as illustrated in the workflow shown in **Fig 1A**. **Table 3** provides a detailed summary of the evaluation metrics for the concatenation method utilizing Repeated Nested Stratified cross-validation. The SVM-RBF model consistently displayed superior performance across all criteria, achieving higher values than SVM-Linear and SVM-Poly, except for sensitivity and specificity. SVM-Poly exhibited heightened sensitivity but marginally lower specificity compared to SVM-RBF and SVM-Linear.

**Fig 1:**
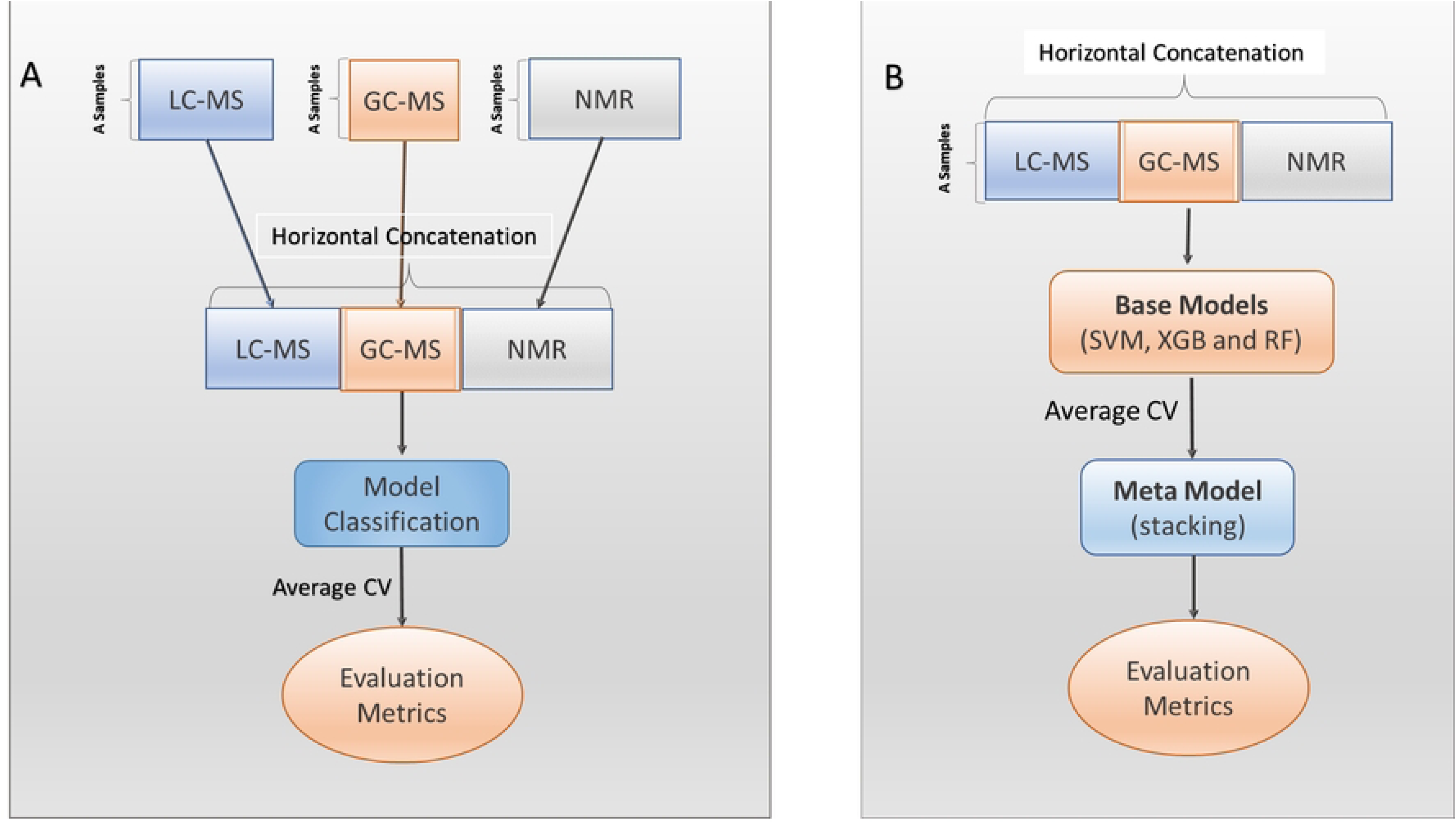
An overview workflow for Straightforward Concatenation and Concatenation-Ensembled approaches. **A)** In the straightforward concatenation method, data is first concatenated and then subjected to nested-stratified cross-validation, followed by the evaluation of SVM models. **B)** In the concatenation-ensemble method, a base model comprising SVM-poly, Random Forest (RF), and XGBoost (XGB) was employed. The base model was trained on the dataset and used to generate predictions for the validation dataset using nested-stratified cross-validation. These predictions from the validation set were subsequently used to train the meta-model.

**Table 3:** Evaluation Metrics for Test Classification of Integrated Metabolomic Platforms. Methods compared include straightforward concatenation with SVMs, concatenation-ensemble with SVMs, RF, and XGB, deep-forest with RF, multiple kernel learning with SVMs, and deep transfer learning with ANNs (scratch and pre-trained). Metrics evaluated include Accuracy (Acc), F1 Score (F1), AUC Score (AUC), Sensitivity (Sen), Specificity (Spe), Balanced Accuracy (BA), and Matthews Correlation Coefficient (MCC).

|  | Acc | BA | F1 Score | AUC | Sen | Spe | MCC |
| --- | --- | --- | --- | --- | --- | --- | --- |
| **Straightforward Concatenation** |  |  |  |  |  |  |  |
| **SVM-Linear** | 0.9540 | 0.9538 | 0.9524 | 0.9338 | 0.9302 | 0.9773 | 0.9089 |
| **SVM-RBF** | 0.9540 | 0.9540 | 0.9535 | 0.9540 | 0.9535 | 95.45 | 0.9080 |
| **SVM-Poly** | 0.9418 | 0.9418 | 0.9434 | 0.9418 | 0.9650 | 0.9186 | 0.8858 |
| **Concatenation- Ensemble** |  |  |  |  |  |  |  |
| **SVM-Poly, RF and XGB** | 0.9770 | 0.9769 | 0.9756 | 0.9769 | 0.9756 | 0.9783 | 0.9539 |
| **Deep - Forest** |  |  |  |  |  |  |  |
| **Cascade-Forest** | 0.9769 | 0.9769 | 0.9769 | 0.9769 | 0.9692 | 0.9846 | 0.9540 |
| **Multiple Kernel Learning** |  |  |  |  |  |  |  |
| **SVM-Linear** | 0.9846 | 0.9846 | 0.9848 | 0.9988 | 1.0000 | 0.9692 | 0.9697 |
| **SVM-RBF** | 0.9923 | 0.9923 | 0.9924 | 0.9998 | 1.0000 | 0.9846 | 0.9847 |
| **SVM-Poly** | 0.9923 | 0.9923 | 0.9924 | 1.0000 | 1.0000 | 0.9846 | 0.9847 |
| **Deep Transfer Learning** |  |  |  |  |  |  |  |
| **Scratch-ANN** | 0.9846 | 0.9855 | 0.9839 | 0.9998 | 1.0000 | 0.9710 | 0.9696 |
| **Pre-Trained-ANN** | 0.9923 | 0.9928 | 0.9919 | 0.9995 | 1.0000 | 0.9855 | 0.9847 |

#### 3-3-2 Concatenation – Ensembled

The ensemble mode for the concatenated dataset involved the utilization of SVM-Poly, Random Forest (RF), and XGBoost as base models. Hyperparameter tuning was conducted through Grid search, with Logistic Regression as the meta-model, and Stacking and Bagging classifiers employed for model training. To address overfitting concerns, Repeated Nested Stratified Cross-Validation was implemented. The comprehensive workflow of Concatenation-Ensembled is illustrated in **Fig 1B**, while **Table 3** presents the improved evaluation metrics attained by Concatenation-Ensembled compared to simple concatenation.

#### 3-3-3 Deep Forest

The workflow of the Deep Forest model is illustrated in **Fig 2**. The Cascade Deep Forest model has a hierarchical structure with multiple layers, where each layer integrates both Random Forest and Extra Trees classifiers to enhance the model’s performance. Initially, the original dataset— comprising concatenated multimodal metabolomics data with prior feature selection—is processed by several forest models in the first layer, producing class probability predictions.

**Fig 2:**
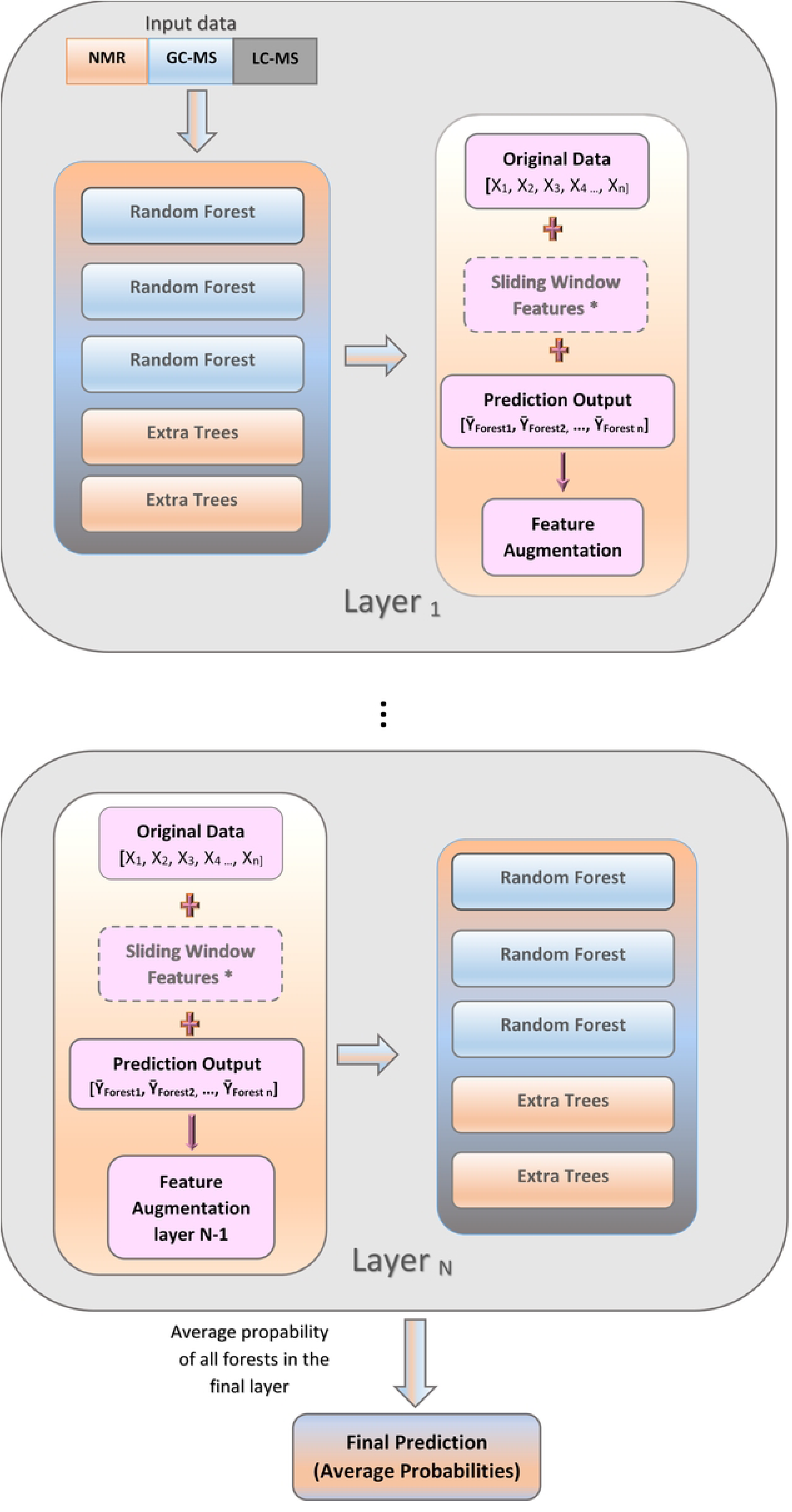
Cascade Deep Forest Strategy Workflow. Concatenated multimodal datasets are initially used as input. Random Forests and Extra Trees independently generate probabilistic predictions at each cascade level. Sliding window features augment the original data, refined through averaging at each level. The final prediction integrates probabilities using an average probabilities strategy across all cascade layers. Note: Sliding window features increased overfitting and were excluded in the final model.

Each layer employs a scanning approach using sliding windows of varying sizes (multi-grained) to extract features from the data. The predictions from the forest models and sliding window features are then integrated with the original features to enrich the feature set. In subsequent layers, this augmented dataset—comprising original features, sliding window features, and predictions from the previous layer—serves as input for new forest models. This iterative process continues layer by layer, enabling the model to capture increasingly complex patterns. In the final layer, the model aggregates the probability predictions from all forest models to derive the final classification decision. This approach effectively leverages the deep, layered structure to enhance predictive accuracy and robustness.

The study identified that using a multi-grained approach resulted in higher overfitting risks, leading to lower evaluation metrics. Figure 2 depicts these findings, with the multi-grained approach shown in a subdued manner. **Table 3** provides a detailed illustration of the evaluation metrics for the Deep Forest model.

The Deep Forest (Cascade-Forest) model achieved balanced and high metrics (Accuracy = 0.9769, AUC = 0.9769), with slightly higher specificity (0.9846) but lower sensitivity (0.9692) compared to the Concatenation-Ensemble model, which had a sensitivity of 0.9756 and specificity of 0.9783. Both models perform similarly, but Deep Forest slightly favors specificity, while the ensemble approach is more balanced in sensitivity and specificity.

#### 3-3-4 Multiple Kernel Learning

MKL was implemented using the EasyMKL algorithm, a scalable approach designed to optimize combination parameters for a predefined set of kernel matrices. **Fig 3A** outlines the primary steps involved in MKL models, and **Fig 3B** provides a schematic illustration of the generation of the Multiview kernel matrix. **Table 3** presents the evaluation metrics for MKL applied to integrating cross-platform LC-MS, GC-MS, and NMR data. Notably, all models, including SVM-Linear, SVM-RBF, and SVM-Poly, demonstrated robust performance, underscoring the efficacy of multiple kernel learning. Furthermore, SVM-RBF and SVM-Poly exhibited slightly higher values for specific metrics than SVM-Linear, suggesting that the non-linear RBF and Polynomial kernels may better capture underlying patterns within the datasets.

**Fig 3:**
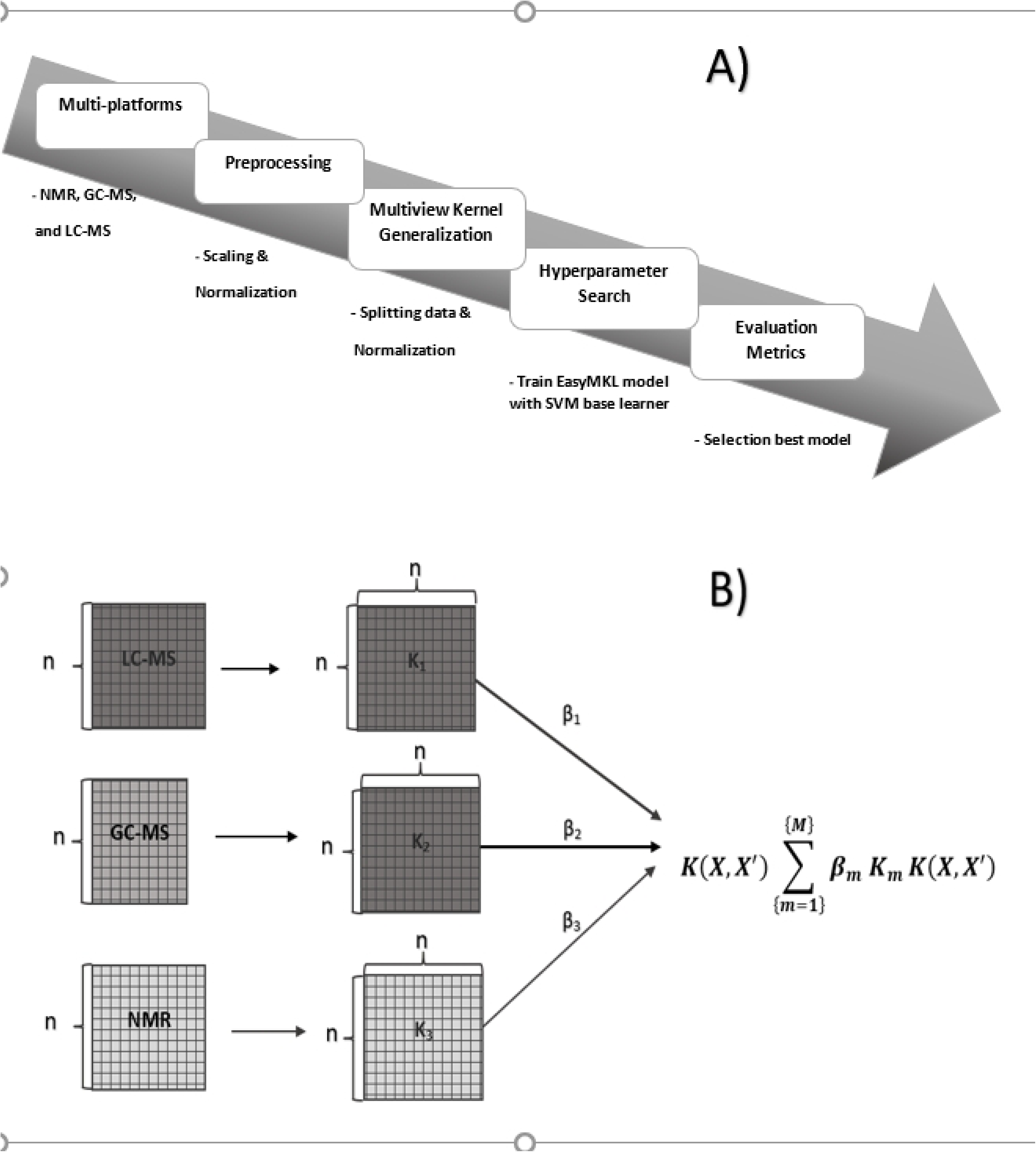
The Multiple Kernel Learning Process. **A)** The figure outlines key stages, including the transition from individual platforms to MKL creation, hyperparameter tuning, and subsequent model evaluation. **B)** The integration of three metabolomics platform datasets into kernel matrices is depicted. These matrices are then combined, incorporating corresponding weights (βm), representing a linear combination of a set of n kernels (K).

#### 3-3-5 Autoencoder Transfer Learning, Extract Feature Weights, and Feature Mapping

The architecture of the custom autoencoder is illustrated in **Fig 4**, highlighting the cross-platform feature concatenation process and the extraction of significant features. Input data from the three distinct platforms were compressed into a compact latent space through an efficient encoding process. The input layer progressed sequentially to a hidden layer with a designated encoding dimension, which played a pivotal role in defining the size of the metabolic profile and effectively reducing the dimensionality of input data from the LC-MS, GC-MS, and NMR platforms. Features from all three platforms were horizontally concatenated to form a multiview dataset for subsequent classification.

**Fig 4.**
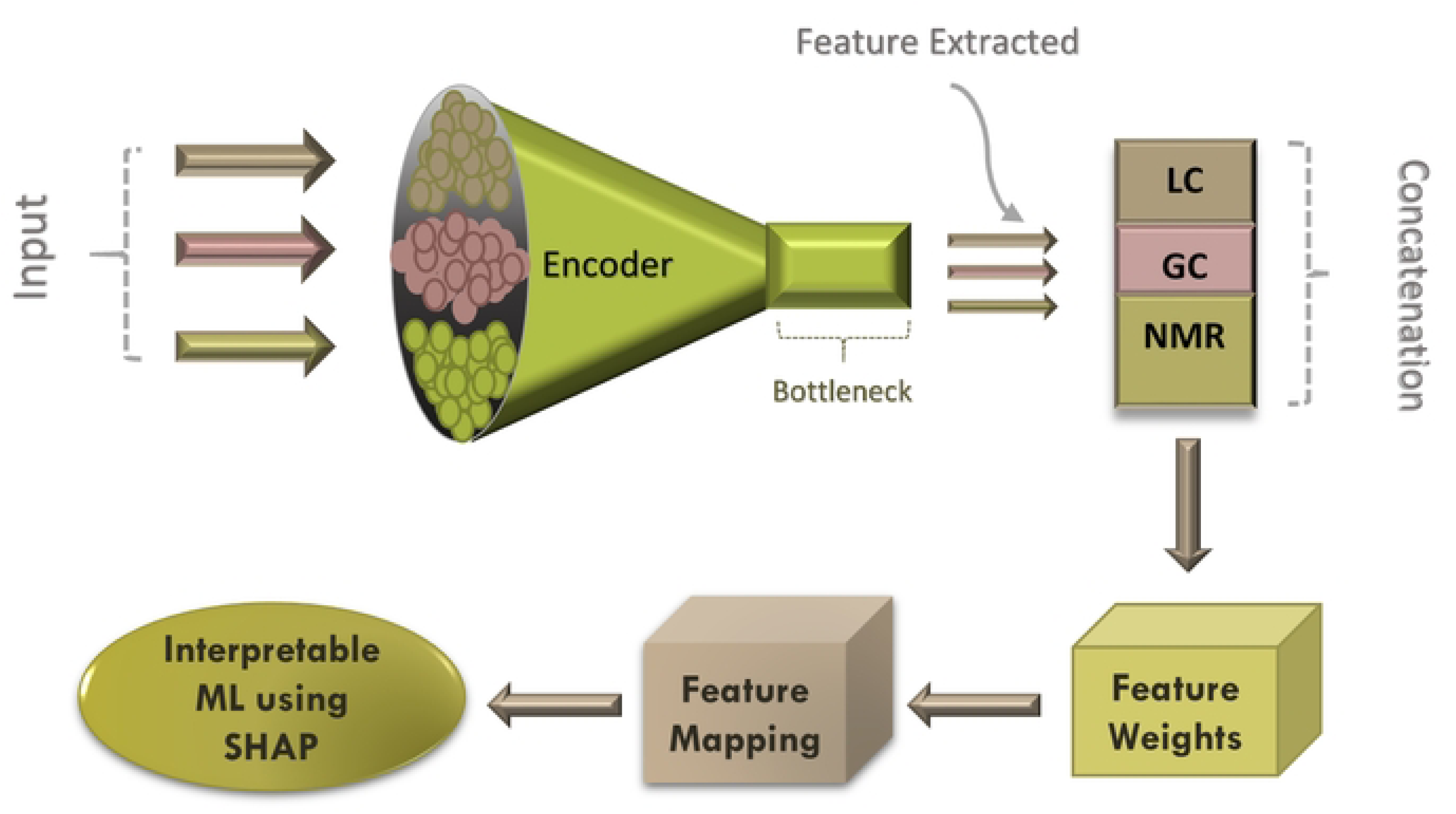
Custom Autoencoder Transfer Learning Architecture. The model utilizes three datasets as input, which are processed through an encoder and a hidden layer for dimensionality reduction. Extracted features from individual platforms are either concatenated and used in subsequent classification steps to identify cancer subtypes or assigned weights and subsequently mapped to identify metabolite names and select key features (30 metabolites per platform). Finally, the top 15 most contributory and abundant metabolites are analyzed using the SHAP method.

In parallel, features extracted from the bottleneck of the encoder for each platform were assigned feature weights. These weights facilitated the identification of significant metabolites, with the top 30 metabolites selected from each platform based on their respective weights. To identify specific metabolite names, features were mapped to detect key metabolites according to their weights. The SHapley Additive exPlanations (SHAP) method was subsequently applied to evaluate the contributions and abundance of these selected metabolites, offering insights into their relevance in the classification process. Furthermore, a Pre-Trained Autoencoder was employed to enhance classification performance and enable more precise subtype detection. This approach utilized optimized neural network parameters from prior training, improving computational efficiency and accuracy by initializing the model with learned representations tailored to all platform datasets.

#### 3-3-6 Interpretable Autoencoder Feature Extraction using SHapley Additive exPlanations (SHAP)

SHAP analysis was conducted using a random forest model on 30 significant metabolites identified across each platform to assess the absolute contributions and abundances of the top 15 metabolites. To ensure a more focused metabolite analysis for the HER2 subtype, ER- or PR-positive samples were excluded, and only HER2-enriched samples were classified as HER2-positive. **S2 Fig** depicts the distribution of these top features, and **Fig 5** presents a comparative analysis of absolute contributions across the three datasets. Within the LC-MS platform, distinct differences were observed between HER2+ and HER2-samples, with mono-unsaturated phospholipids such as PC 30:1, PE 32:1, and SM 32:1 showing elevated levels in HER2+ samples. Triacylglycerols (TAGs) displayed varying patterns: TAG 45:1, TAG 48:2, and TAG 55:3 were reduced in HER2+ samples, while TAG 52:5, TAG 55:2, TAG 58:10, and TAG 60:2 showed increased levels. In the GC-MS platform, higher contributions were observed in HER2+ samples for beta-alanine, glycine, oleic acid, gluconic acid, cysteine, and lysine, whereas 5’-deoxy-5’-methylthioadenosine (MTA) and nicotinamide had more significant contributions in HER2-samples. The NMR analysis indicated elevated levels of glutamate, methionine, cholate, N-acetyl aspartic acid, aspartic acid, beta-alanine, and lysine in HER2+ samples. In contrast, isoleucine, leucine, o-phosphocholine, glycerophosphocholine, glutamine, and myoinositol demonstrated increased contributions in HER2-samples. **Fig 6** illustrates the comparative abundance of the top 15 metabolites across the individual platforms. In HER2-positive samples, the following metabolites demonstrated increased levels: PC 30:1, PE 32:1, SM 32:1, SM 36:1, TAG 55:2, TAG 55:3, TAG 52:4, TAG 52:5, TAG 50:4, gluconic acid, beta-alanine, oleic acid, cysteine, tyrosine, and pyrogallol. In contrast, nicotinamide, arabinose, palmitic acid, 5’-deoxy-5’-methylthioadenosine (MTA), Cer 42:2, cytidine-5-diphosphate, and TAG 48:2 exhibited increased abundance in HER2-negative samples. This study emphasizes machine learning biomarker discovery. Meanwhile, metabolites were selected based on statistical analysis criteria, precisely an effect size (Cohen’s d) greater than 0.5 (absolute value) and a p-value from a Student’s t-test of less than 0.05 (see **Table 4**). The selected metabolites include PC 30:1, PE 32:1, SM 32:1, β-alanine, MTA, and N-Acetylaspartic acid.

**Fig 5:**
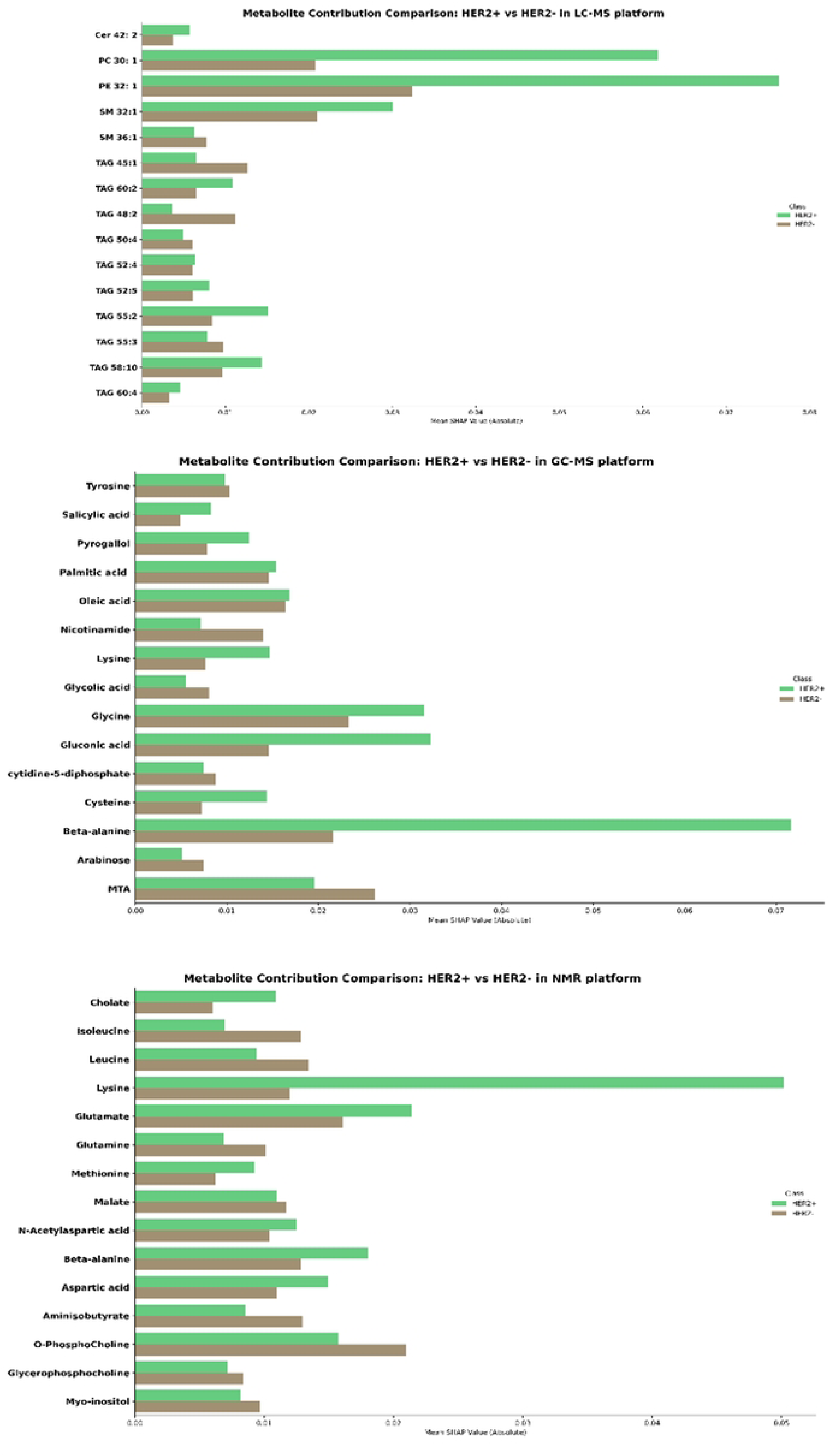
Comparison of the top 15 features contributing to the classification of HER2+ and HER2-breast cancer subtypes based on SHAP values. Each bar represents a given metabolite’s mean absolute SHAP value, indicating its relative importance in distinguishing between the two subtypes. The different colors correspond to the HER2 status: **dark blue** for HER2-positive and **green ocean** for HER2-negative.

**Fig 6:**
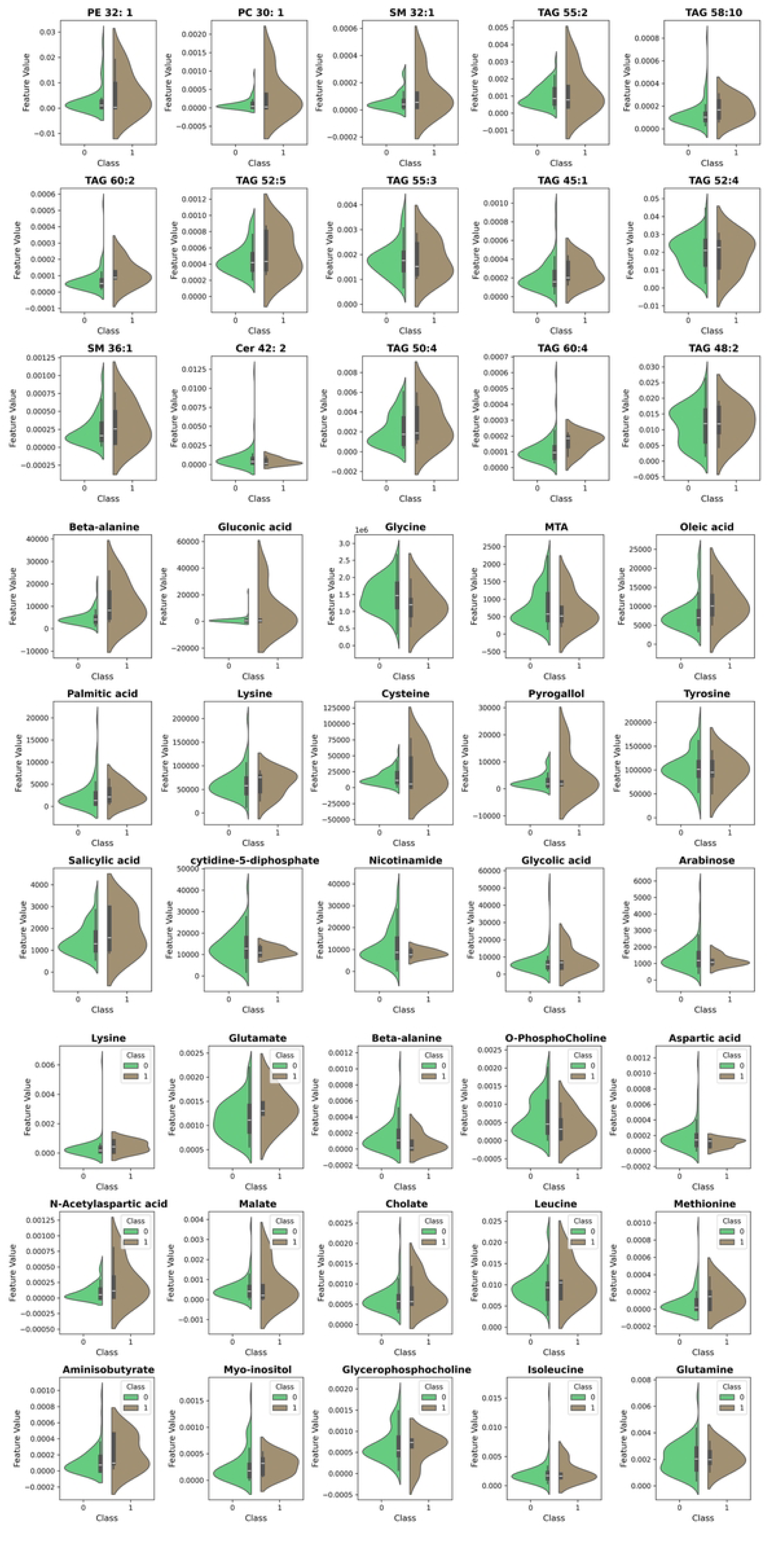
Violin plots representing the abundance comparison of the top 15 metabolites for the classification of HER2+ and HER2-breast cancer subtypes across three platforms: LC-MS, GC-MS, and NMR. The analysis is based on SHAP values. The width of each violin represents the density of data points at various feature values, providing a visual comparison of metabolite abundance between the two subtypes.

**Table 4:** Statistical selection of metabolite profiles based on an effect size threshold of |0.5| and a p-value < 0.05 for HER2-positive versus HER2-negative cases.

| No | Metabolite | Effect-size | P-value |
| --- | --- | --- | --- |
| 1 | **PC 30: 1** | 1.446674 | 0.011975 |
| 2 | **PE 32: 1** | 0.948638 | 0.015945 |
| 3 | **SM 32:1** | 0.859051 | 0.035398 |
| 4 | **Beta-alanine** | 0.651673 | 0.001516 |
| 5 | **MTA** | 0.511539 | 0.013456 |
| 6 | **N-Acetylaspartic acid** | 0.856065 | 0.019358 |

#### 2-3-6 Classification Using Artificial Neural Network

**Fig 7** depicts the schematic workflow of the sequential Artificial Neural Network (ANN) model used to analyze datasets. The model consists of a dense layer with 128 nodes following an input layer that incorporates datasets from three different platforms.

**Fig 7:**
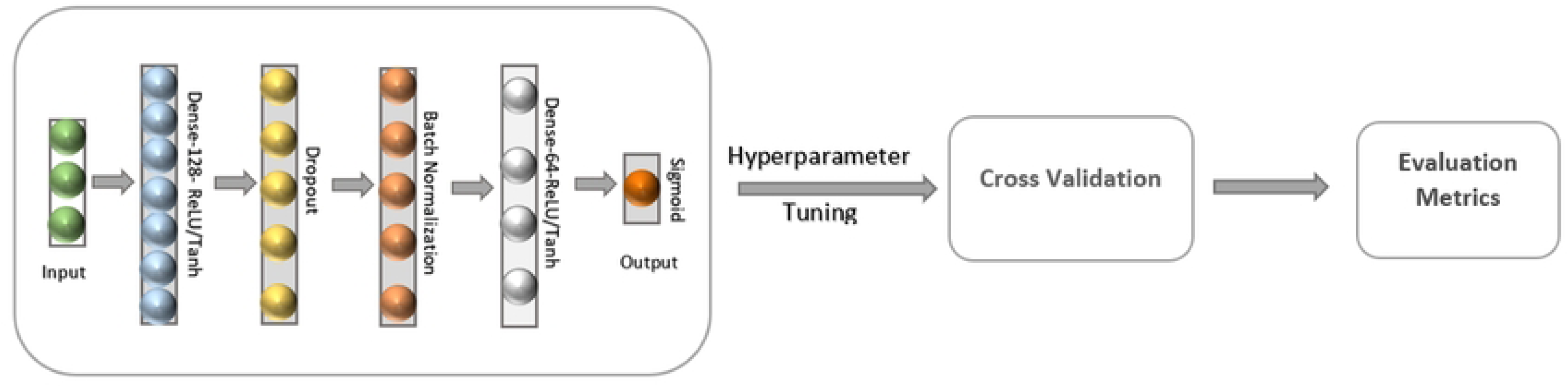
The architecture of the ANN designed for concatenated data. The subsequent results underwent hyperparameter tuning and cross-validation using a repeated 5-fold stratified cross-validation approach.

Subsequently, a dropout layer is introduced to mitigate overfitting and enhance generalization, followed by batch normalization, a critical technique for stabilizing and accelerating the training process. A dense layer with 64 nodes is added to extract hierarchical features, utilizing Tanh activation functions to capture intricate details within the multimodal data. Finally, the output layer is tailored for binary classification methods, comprising a single node activated by the sigmoid function. All hyperparameters are fine-tuned through repeated stratified cross-validation.

In **Table 3**, the ANN models exhibited exceptional performance across various metrics. A comparison of evaluation metrics between the Scratch-ANN and Pre-Trained-ANN models reveals that the Pre-Trained-ANN outperformed the Scratch-ANN in several key areas: Accuracy (0.9923 vs 0.9846), Balanced Accuracy (0.9928 vs 0.9855), F1 Score (0.9919 vs 0.9839), and MCC (0.9847 vs 0.9696). Notably, both models achieved high AUC values (0.9995 for Pre-Trained-ANN and 0.9998 for Scratch-ANN), perfect Sensitivity (1.0000), and substantial Specificity (0.9855 for Pre-Trained-ANN and 0.9710 for Scratch-ANN).

#### 2-3-7 A Comprehensive Comparison of Integration Multimodal

A comprehensive comparison of Accuracy, F1 Score, and MCC across integrated multimodal approaches and individual platforms is presented in **Fig 8A**. The top-performing models for each method were selected to facilitate a comparison of the best performances across various platforms. The Deep Transfer Learning and MKL approaches, specifically SVM-RBF and SVM-Poly, were identified as the leading models, exhibiting outstanding performance in accuracy, F1 Score, and MCC. The Concatenated Ensemble and Deep Forest ranked second and demonstrated strong performance across all metrics, confirming their positions as competitive and integrated multimodal solutions. Individual platforms, particularly LC-MS, displayed impressive accuracy and F1 Scores. The Concatenated platform also achieved competitive results, reflecting average performance across LC-MS, GC, and NMR datasets. **Fig 8B** illustrates the learning curves for accuracy and loss values during the training and cross-validation phases. In contrast, **Fig 8C** presents the evaluation metrics, including accuracy, F1 score, AUC, balanced accuracy, and MCC, within the framework of transfer deep learning. The strong consistency observed between the learning curves for training and cross-validation and among the evaluation metrics for training, testing, and cross-validation suggests that a robust predictive model has been developed. **Fig 8D** depicts the learning curves for MKL, demonstrating high consistency between the training and cross-validation results. In contrast, **Fig 8E** presents the evaluation metric plots, revealing that cross-validation performs at a lower level than both training and testing, although the latter two demonstrate strong consistency.

**Figure 8:**
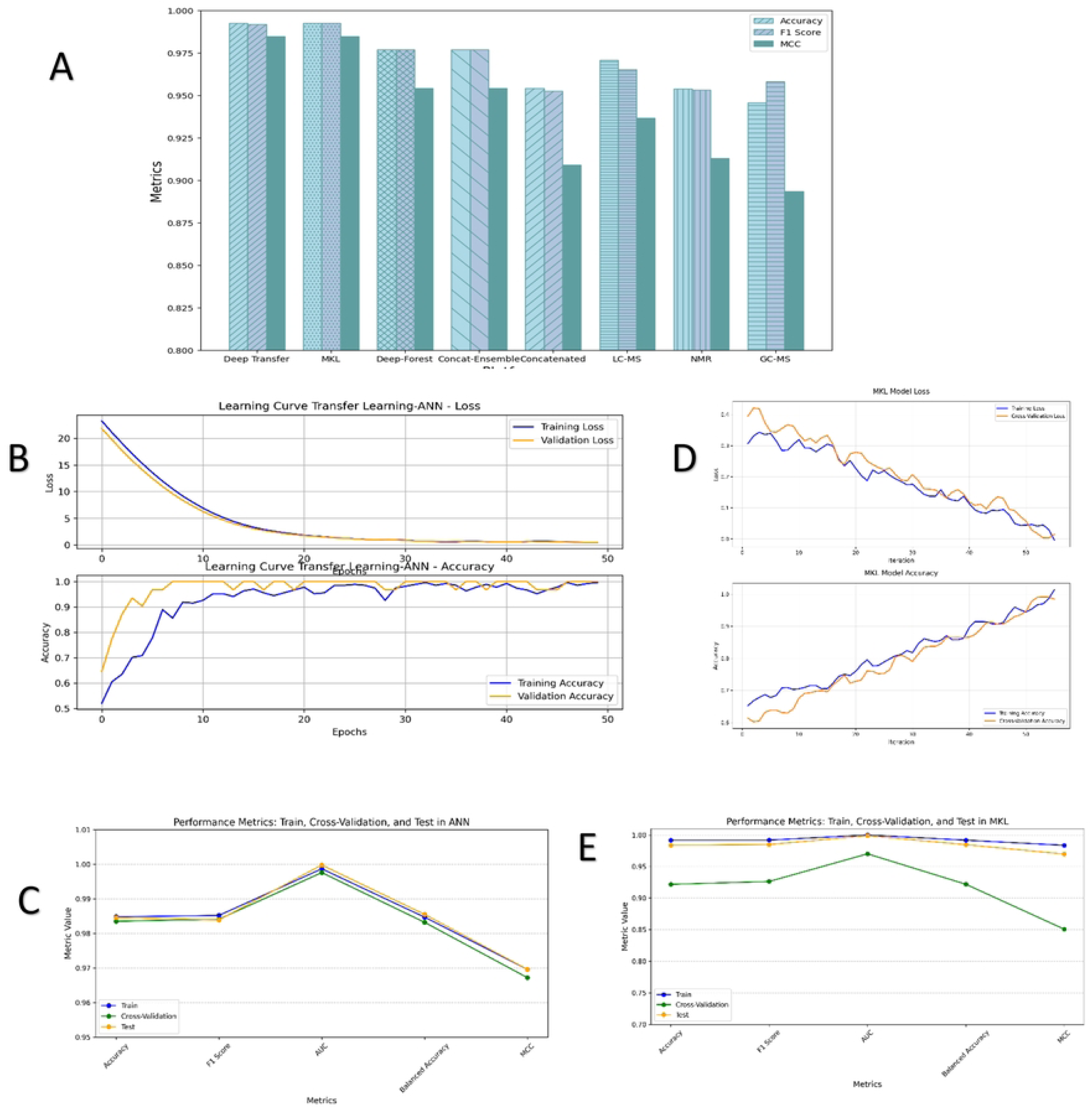
Performance Evaluation of Predictive Models in Multimodal and Individual Platforms. **A)** A comparison of accuracy, F1 score, and MCC for integrated multimodal approaches versus individual platforms reveals that the top-performing models, including Deep Transfer Learning and MKL approaches (SVM-RBF, SVM-Poly), stand out for their exceptional performance across these metrics. **B)** Learning curves illustrating accuracy and loss values during the training and cross-validation phases in transfer deep learning. **C)** Evaluation metrics, including accuracy, F1 score, AUC, balanced accuracy, and MCC, are presented within the transfer deep learning framework. These metrics demonstrate the model’s performance across training, testing, and cross-validation phases. **D)** Learning curves for MKL approaches demonstrate high consistency between training and cross-validation results. **E)** The plot of evaluation metrics reveals that cross-validation performance is lower than training and testing, which suggests potential issues related to data variability, model complexity, or overfitting. Key: Concat=Concatenation.

#### 2-3-8 Average and Grand effect size of the individual platform

**Table 5** presents the average and grand effect sizes for each platform. Notably, all platforms demonstrate effect sizes exceeding 0.5, with accuracies surpassing 80%. This finding aligns with prior research suggesting that a small to moderate sample size can be sufficient for analysis when using high-quality datasets. Among the platforms, GC-MS stands out with significantly higher average and grand effect sizes (greater than 2) than the others.

**Table 5:**
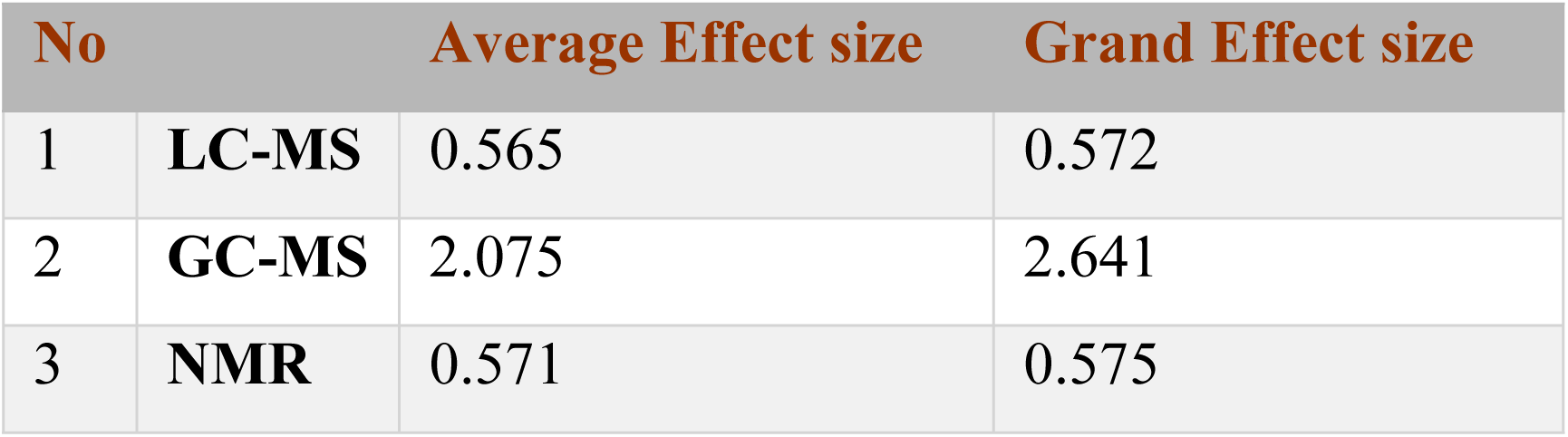
Average and Grand effect size of individual platforms.

## 4 Discussion

In the present study, the optimal feature selection method was meticulously chosen based on the superior performance demonstrated by the SVM-linear technique across four specified methods (ReliefF, SVM-RFE, RF-RFE, and Mutual Information). Notably, SVM-RFE exhibited exceptional performance in LC-MS and NMR datasets, while comparable performances were observed among all methods, except RF-RFE, in GC-MS data. To ensure simplicity and consistency, we standardized the feature selection approach across all platforms, selecting SVM-RFE as the preferred method. Before initiating comparisons, we carefully fine-tuned the optimal number of features for each method on individual platforms.

Transfer deep learning was implemented using a custom autoencoder that utilized three metabolomics modalities as input, with dimensionality reduction performed in the hidden layer of the encoder as output. Features extracted from the encoder’s bottleneck were analyzed using feature weights, generating feature maps identifying reduced-dimensional features. The top 30 metabolites from each analytical platform were selected based on this analysis. Subsequent SHAP analysis refined this selection to the top 15 metabolites per platform. The SHAP analysis, which compared contributions and abundance values between HER2-positive and HER2-negative samples in the LC-MS platform, revealed elevated levels of mono-unsaturated phospholipids— specifically PC 30:1, PE 32:1, and SM 32:1—in HER2-positive samples. This elevation corresponds to increased Stearoyl-CoA Desaturase-1 (SCD1) expression in HER2-positive breast cancer cells, particularly in cases negative for estrogen receptor (ER) and progesterone receptor (PR) (47, 48). SCD1 plays a crucial role in breast cancer by influencing membrane fluidity and promoting cancer cell proliferation and survival. It facilitates the conversion of saturated fatty acids to monounsaturated fatty acids, essential for maintaining membrane integrity and supporting tumor growth (49, 50). Triacylglycerol (TAG) species showed distinct patterns in their contributions and abundances. Specifically, TAG 55:2, TAG 52:5, and TAG 58:10 were elevated in HER2-positive samples, reflecting increased lipid droplet formation and fat storage in cancer cells. Conversely, TAG 48:2, TAG 50:4, and TAG 55:3 levels were decreased in HER2-positive samples, likely due to enhanced TAG consumption during tumor cell division and proliferation. In GC-MS analysis, palmitic acid (C16:1) and oleic acid (C18:1) showed increased contributions and abundances due to enhanced SCD1 activity. These fatty acids, synthesized via SCD1-mediated pathways, are crucial precursors for mono-unsaturated phospholipids such as PC 30:1 and PE 32:1. Elevated gluconic acid levels have been identified in HER2-positive breast cancer. As a byproduct of glucose oxidation, gluconic acid plays a pivotal role in cellular metabolism and is frequently dysregulated in cancer cells. Increased levels of gluconic acid are linked to altered glycolysis and heightened oxidative stress in breast cancer models. Furthermore, gluconic acid can serve as an alternative carbon source, providing cancer cells with metabolic flexibility that supports their survival and proliferation. In the context of breast cancer, this elevation in gluconic acid may indicate the metabolic reprogramming that underlies cancer development and progression (51, 52). In addition, the GC-MS dataset reveals a significant increase in the contribution and abundance of β-alanine in HER2-positive breast cancer. Elevated levels of β-alanine may reflect a compensatory endogenous mechanism that counteracts tumor-induced oxidative stress and sustains tumor growth through aberrant metabolic pathways. This finding underscores its involvement in the metabolic reprogramming of amino acids, which supports the enhanced glucose utilization and energy metabolism required for rapid cancer cell proliferation (53). Interestingly, a plausible hypothesis suggests that β-alanine’s potential as an anticancer agent in breast cancer is multifaceted, depending on whether it is endogenous or exogenous. Elevated endogenous β-alanine may aid tumor survival by facilitating metabolic reprogramming and managing oxidative stress. In contrast, exogenous β-alanine supplementation could shift this balance toward anticancer effects by increasing carnosine levels beyond the tumor’s metabolic control. This increase may disrupt glycolysis, buffer intracellular acidity, and selectively amplify oxidative stress in cancer cells while preserving normal tissue function. Furthermore, carnosine’s ability to modulate critical oncogenic pathways, such as PI3K/AKT/mTOR, along with its potential for epigenetic regulation, could further impair tumor progression and enhance the efficacy of immune-based therapies(54–57). These combined effects may collectively disrupt the metabolic demands and invasive capacity of HER2-positive tumor cells, positioning exogenous β-alanine as a promising therapeutic adjunct in this aggressive cancer subtype. Nevertheless, further studies are needed to analyze the effects of β-alanine as a supplement or therapeutic agent in this context.

Nicotinamide levels and contributions are significantly decreased in HER2-positive breast cancer. Previous studies suggest that nicotinamide, combined with non-steroidal anti-inflammatory drugs (NSAIDs), can bind to and inhibit the SIRT1 deacetylase. This enzymatic inhibition activates the anti-proliferative p53/p21 pathway, reducing the risk of invasive breast cancer subtypes, including HER2-positive and triple-negative cases(58, 59). The observed low levels of nicotinamide in HER2-positive breast cancer may serve as a potential therapeutic biomarker, guiding strategies to mitigate the aggressiveness of this cancer subtype.

In addition, 5’-methylthioadenosine (MTA) shows reduced contribution and abundance in HER2-positive tumors compared to HER2-negative tumors. MTA has demonstrated potential as an anti-tumor agent by inhibiting tumor cell proliferation and invasion while promoting apoptosis. These effects are linked to MTA’s ability to regulate the inflammatory microenvironment of tumors, potentially creating conditions that are less conducive to cancer progression (60, 61). Furthermore, in the NMR platform, elevated levels of lysine, N-acetylaspartic acid, O-phosphocholine, and cholate were observed in HER2-positive breast cancer. The increase in lysine and glutamate are particularly associated with the reprogramming of amino acid synthesis, and N-acetyl aspartic acid, O-phosphocholine, and cholate are linked to promoting tumor growth, metastasis, cell proliferation, and enhanced cancer survival (62–64). Based on a defined effect size and p-value threshold, statistical analysis identified a subset of six metabolites—PC 30:1, PE 32:1, SM 32:1, β-alanine, MTA, and N-acetylaspartic acid—across all platforms. While these metabolites underscore the significance of these biomarkers, further investigation using SHAP analysis is prioritized to identify additional biomarker signatures.

Evaluation of individual binary classifications within the HER2 group applied SVM-RFE feature selection for SVM models, revealing distinct performance characteristics. To improve robustness, hyperparameter tuning was conducted using grid search, repeated stratified cross-validation, and L2 regularization, effectively mitigating overfitting and enhancing generalization. In NMR datasets, the SVM-Linear model exhibited higher performance, achieving notable accuracy, sensitivity, and MCC values. Conversely, in GC-MS data, SVM-RBF excelled in accuracy and F1 score, while SVM-Linear demonstrated superiority in sensitivity and AUC. Across all platforms, LC-MS emerged as the top performer, achieving remarkable accuracy, F1 score, sensitivity, and MCC through SVM polynomial.

Straightforward concatenation resulted in accuracy, F1 score, AUC, and specificity that were either higher or equivalent to those observed in the GC-MS and NMR datasets. However, compared to LC-MS, the performance was considered average. The Concatenation-Ensemble and Deep-Forest methods consistently outperformed the individual datasets across all metrics. Employing Easy-MKL for SVM models in Multiple Kernel Learning demonstrated outstanding performance, highlighting robust capabilities in classification tasks. Specifically, SVM-RBF and SVM-Poly demonstrated remarkably high accuracy, F1 score, AUC, and MCC. Additionally, SVM-Linear displayed robust overall performance, particularly excelling in sensitivity and specificity. These findings support previous studies, indicating enhanced performances of multimodality compared to individual platforms(22, 26). In contrast, the Multimodal Deep Learning approach, which used a custom autoencoder architecture to extract features from individual platforms within the hidden layer of the encoder and then concatenated these features horizontally for input into the ANN classifier, demonstrated outstanding performance. Furthermore, using a pre-trained autoencoder potentially enhanced performance by starting from a well-optimized state rather than from an untrained state, which had been shown to yield slightly higher results.

The comprehensive comparison of integrated multiview metabolite performance revealed that MKL and Deep Transfer Learning secured the top positions across all performance metrics, providing a well-balanced estimation. The concatenation-ensemble and Deep Forest methods ranked second, exhibiting relatively strong evaluation metrics. The analysis of learning curves for accuracy and loss, along with the evaluation metrics for training, testing, and cross-validation in Deep Transfer Learning, reveals a remarkable level of consistency. On the other hand, MKL exhibits a high level of consistency in learning curves between training and cross-validation. However, comparing performance between training and testing datasets shows a strong correlation, while cross-validation results show lower performance. This inconsistency may be due to various factors, such as overfitting caused by model complexity, potential challenges related to early stopping, and difficulties associated with the high dimensionality of the kernel matrix.

The identification of integration methods requiring optimal computational resources poses significant challenges. While the concatenation ensemble typically relies on simpler operations, such as concatenation and basic ensemble techniques, adopting optimization methods for each model increases computational complexity. In Deep Forest, multiple layers of decision trees are trained, resulting in moderate computational complexity. However, Deep Forest tends to have higher computational complexity and longer running times than concatenated-ensemble methods due to the need to train and evaluate multiple layers of decision trees iteratively. Furthermore, in Concatenation-Ensemble, Deep-Forest, and MKL methods, individual platform features are initially selected before integration. In Deep Transfer, features are specified using an autoencoder, followed by an artificial neural network (ANN) classification, indicating a moderately complex computational process. When employed solely for feature extraction, autoencoders yield a succinct data representation, contributing to reduced complexity compared to more intricate methods. However, achieving generalization and conducting hyperparameter tuning can incur substantial computational expenses. Integrated multimodal analysis has demonstrated outstanding performance in detecting HER2-positive and HER2-negative cases, crucially impacting sensitive metrics such as sensitivity and specificity to minimize false negatives and false positives, respectively. The robustness of these approaches significantly enhances the accuracy and reliability of determining HER2 status, thereby improving precision outcomes. Achieving sensitivity = 1 in MKL and Deep Transfer Learning may imply a risk of overfitting, mainly due to potential issues like oversampling. However, there is typically high consistency between the training, testing, and cross-validation phases. Implementing techniques such as oversampling and repeated stratified cross-validation mitigates these risks, ensuring a more robust model evaluation and deployment.

Due to the relatively small size of each platform dataset (423 samples for individual datasets after oversampling), which is often considered insufficient for Deep Learning methodologies, this study acknowledges a recognized phenomenon observed in prior research where ML accuracies tend to stabilize after a certain number of samples in sufficiently high-quality data. This study evaluated dataset quality by considering average and grand effect sizes of data, focusing specifically on those with effect sizes greater than 0.5 and accuracies exceeding 80% for robust classification(14). Moreover, while larger sample sizes are typically beneficial in high-dimensional data analysis, such as genomics or imaging, where they aid in reducing the likelihood of overfitting, metabolomics may offer the opportunity for practical analysis with smaller sample sizes. This can be attributed to the intrinsic nature of metabolomics data, which captures the end products of cellular processes, in addition to the advantages of precise feature selection and the biological significance of metabolites.

Integrated multimodal metabolomics using MKL has demonstrated considerable promise, particularly in cancer detection and the identification of molecular subtypes. However, it encounters specific challenges in feature analysis, limiting its suitability for detecting distinct biomarkers. MKL integrates composite features from diverse platforms, complicating the analysis of metabolite signatures within the transformed feature space. To address these limitations, feature extraction from individual platforms using autoencoders, followed by analysis with the SHAP technique, has proven effective. This approach emphasizes the importance of deep transfer learning as both an advanced classification module and a tool for identifying biomarker signatures. Nonetheless, a significant challenge lies in obtaining corresponding data across multiple metabolomics platforms for the same set of samples, which limits the scope of this research. Additionally, the absence of external validation presents a critical drawback, as independent validation is essential for confirming findings and ensuring broader applicability. Despite these challenges, this study establishes a synergistic pipeline for integrated multimodal metabolomics analysis. The Deep-Transfer strategy facilitates robust and scalable integration of high-dimensional biological data across various omics layers, enabling reliable biomarker discovery.

By addressing the complexities inherent in multi-omics integration, this methodology significantly enhances cancer detection, precise classification of cancer subtypes, and biomarker identification. Furthermore, this approach holds substantial potential for advancing translational oncology research. It could improve outcomes in difficult-to-treat cancers, such as pancreatic and hepatocellular carcinoma, by enabling early detection, enhancing predictive accuracy, and uncovering clinically relevant biomarker profiles. The comprehensive framework presented in this study sets the stage for broader applications in multi-omics and paves the way for transformative advancements in cancer research.

### Data availability statement

The research data and code are readily accessible on CodeOcean, a platform designed for transparent and reproducible research. The Compute Capsule’s provisional DOI for the code is 10.24433/CO.3436717.v5, ensuring easy retrieval and referencing for interested parties.

